# FMNL2 regulates actin for ER and mitochondria distribution in oocyte meiosis

**DOI:** 10.1101/2023.10.05.561058

**Authors:** Meng-Hao Pan, Zhen-Nan Pan, Ming-Hong Sun, Xiao-Han Li, Jia-Qian Ju, Shi-Ming Luo, Xiang-Hong Ou, Shao-Chen Sun

## Abstract

During mammalian oocyte meiosis, spindle migration and asymmetric cytokinesis are unique steps for the successful polar body extrusion. The asymmetry defects of oocytes will lead to the failure of fertilization and embryo implantation. In present study we reported that an actin nucleating factor formin-like 2 (FMNL2) played critical roles in the regulation of spindle migration and organelle distribution. Our results showed that FMNL2 mainly localized at the oocyte cortex and periphery of spindle. Depletion of FMNL2 led to the failure of polar body extrusion and large polar bodies in oocytes. Live-cell imaging revealed that the spindle failed to migrate to the oocyte cortex, which caused polar body formation defects, and this might be due to the decreased polymerization of cytoplasmic actin by FMNL2 depletion. Furthermore, mass spectrometry analysis indicated that FMNL2 was associated with mitochondria and endoplasmic reticulum-related proteins, and FMNL2 depletion disrupted the function and distribution of mitochondria and endoplasmic reticulum, showing with decreased mitochondrial membrane potential and the occurrence of endoplasmic reticulum stress. Microinjecting Fmnl2-EGFP mRNA into FMNL2-depleted oocytes significantly rescued these defects. Thus, our results indicate that FMNL2 is essential for the actin assembly, which further involves into meiotic spindle migration and ER/mitochondria functions in mouse oocytes.

## Introduction

Mammalian oocyte maturation is an asymmetric division process that generates a large egg and a small polar body. This asymmetry is critical for the following fertilization and early embryo development. After germinal vesicle breakdown (GVBD), the meiotic spindle is organized at the center of the oocyte, and then it migrates to the oocyte cortex at the late metaphase I (MI). The oocytes are arrested at metaphase II (MII) after the extrusion of first polar body (*1, 2*). Actin filaments, as the most widely distributed cytoskeleton in cells, regulate various dynamic events during oocyte meiotic maturation (*3*), and two key events are the spindle migration and cortical reorganization in mammalian oocytes (*1, 4, 5*). Small GTPases and actin nucleation factors are shown to promote the assembly and function of actin. The actin nucleation factors are the molecules that directly promote the actin assembly: Arp2/3 complex control the assembly of branched actin, and formin family member Formin2 (FMN2) and Spire1/2 control the assembly of linear actin. These proteins are all proposed to play a role in actin-related spindle migration and cytokinesis during mammalian oocyte maturation (*6–8*). The cortex protein Arp2/3 complex nucleates the actin to produce a hydrodynamic force to move the spindle toward the cortex, and regulates cytokinesis during oocyte maturation (*1, 8*). FMN2 and Spire1/2 nucleates actin around the spindle in the cytoplasm to give the meiotic spindle an initial power for migration (*7, 9*).

Besides Formin2, the DRFs (diaphanous-related formins) subfamily in the formin family has been extensively studied. The DRFs family consists of mDia, Daam, FHOD and FMNLs (*10*). The “Formin-like” proteins (FMNLs) subfamily includes FMNL1 (FRL1), FMNL2 (FRL3), and FMNL3 (FRL2). Like other Formin family proteins, FMNLs play important roles in cell migration, cell division, and cell polarity (*10, 11*). While FMNL2 is widely expressed in multiple human tissues, especially in the gastrointestinal and mammary epithelia, lymphatic tissues, placenta, and reproductive tract (*12*). As an important actin assembly factor, FMNL2 accelerates the elongation of actin filaments branched by Arp2/3 complex (*13*). In invasive cells, FMNL2 is mainly localized in the leading edge of the cell, lamellipodia and filopodia tips, to improve cell migration ability by actin-based manner (*13–15*). FMNL2 is also involved in the maintenance of epithelial-mesenchymal transition (EMT) in human colorectal carcinoma cell (*16*). Besides its roles on the actin assembly, emerging evidences indicate that FMNL2 may interact with organelle dynamics. It is shown that FMNL2 is related with the Golgi apparatus, since the absence of FMNL2/3 can cause the Golgi fragmentation (*17*). However, till now the roles of FMNLs especially FMNL2 on oocyte meiosis are still largely unknown.

In the present study, we disturbed the FMNL2 expression and explored the roles of FMNL2 during mouse oocyte meiosis. Our results showed that FMNL2 was essential for the polar body size control and successful extrusion; and these abnormal phenotypes might be due to aberrant actin-based meiotic spindle migration. Meanwhile, we also found that FMNL2 was essential for the functions and distribution of mitochondria and endoplasmic reticulum. Therefore, this study provided the evidence for the critical roles of FMNL2-mediated actin on spindle movement and organelle dynamics in mammalian oocytes.

## Materials and Methods

### Antibodies and chemicals

Rabbit monoclonal anti-FMNL2 antibody, rabbit monoclonal anti-Arp2 antibody, mouse monoclonal anti-profilin1 antibody were from Santa Cruz (Santa Cruz, CA, USA). Rabbit monoclonal anti-Fascin antibody was purchased form Abcam (Cambridge, UK). Rabbit polyclonal anti-INF2 antibody was purchased from Proteintech (Proteintech, CHI, USA). Rabbit monoclonal anti-α-tubulin (11H10) antibody, rabbit monoclonal anti-Grp78 antibody, rabbit monoclonal anti-cofilin antibody and rabbit monoclonal anti-Chop antibody were from Cell Signaling Technology (Beverly, MA, USA). Mouse monoclonal anti-α-tubulin-FITC antibody was from Sigma-Aldrich Corp. (St. Louis, MO, USA). FITC-conjugated goat anti-rabbit IgG were from Zhongshan Golden Bridge Biotechnology (Beijing). ER-Tracker Red, Mito-Tracker Green and enhanced mitochondrial membrane potential assay Kit were from Beyotime Biotechnology (Shanghai). All other chemicals and reagents were from Sigma-Aldrich Corp., unless otherwise stated.

### Ethics statement and oocyte culture

We followed the guidelines of Animal Research Institute Committee of Nanjing Agricultural University to conduct the operations. The animal facility had license authorized by the experimental animal committee of Jiangsu Province (SYXK-Su-20170007). The Female Institute of Cancer Research (ICR) mice, aged 4–6 weeks, were kept in a room with a regulated temperature of 22 °C and provided with a standard diet. Fully developed germinal vesicle stage (GV) oocytes were retrieved from the ovaries of mice, and then cultured in M16 medium with paraln oil at 37 °C and in the presence of 5% CO2 for in vitro maturation. At specific intervals, the oocytes were collected for various tests and analyses. The oocytes were placed at 37°C with an atmosphere of 5% CO2, and cultured to different time points for immunostaining, microinjection and western blot.

### Plasmid construct and in *vitro* transcription

Template RNA was generated from mouse ovaries with RNA Isolation Kit (Thermos fisher), then we reversed transcription of these RNA to create cDNA by a PrimeScript 1st strand cDNA synthesis kit (Takara, Japan). FMNL2-EGFP vector was generated by Wuhan GeneCreate Biological Engineering Co, Ltd. mRNA was synthesized from linearized plasmid using HiScribe T7 high yield RNA synthesis kit (NEB), then capped with m7G (5’) ppp (5’) G (NEB) and tailed with a poly(A) polymerase tailing kit (Epicentre) and purified with RNA clean & concentrator-25 kit (Zymo Research).

### Microinjection of FMNL2 siRNA and mRNA

The FMNL2 siRNA working solution was dissolved in RNase-free water, achieving a concentration of 20 μM. For FMNL2 knockdown (KD), three individual siRNA strands were precisely mixed and subjected to centrifugation to obtain the supernatant, approximately 5-10 pL of supernatant was microinjected into the cytoplasm of GV stage oocytes. In contrast, an equal volume of a negative control solution was microinjected into the cytoplasm of oocytes in the control group. FMNL2 siRNA: 5′-GCU GAA UGC UAU GAA CCU ATT-3′, 5′-GCC AUU GAU CUU UCU UCA ATT-3′, 5′-GGA AUU AAG AAG GCG ACA ATT-3′; Negative control siRNA: 5′-UUC UCC GAA CGU GUC ACG UTT-3′. Then, the oocytes were arrested in the GV stage for 18h in M16 medium with 5μM milrinone. This was done to optimize the effectiveness of the siRNA and aid in the depletion of FMNL2. For the rescue experiment, 5–10 pL of 200ng/μL FMNL2-EGFP mRNA was injected into the GV oocytes 18h after FMNL2 siRNA injection. Following that, the GV stage oocytes were cultured in M16 medium with 5μM milrinone for 4h. Then, the oocytes being cultured in fresh M16 medium for subsequent experiments.

### Immunofluorescent staining and confocal microscopy

Oocytes were fixed in 4% paraformaldehyde (in PBS) for 30 min and permeabilized with 0.5% Triton X-100 in PBS for 20min then blocked in blocking buffer (1% BSA-supplemented PBS) at room for 1 h. For FMNL2 staining, the oocytes at various stages (GV, GVBD, MI, and MII) were incubated with Rabbit monoclonal anti-FMNL2 antibody (1:100) at 4 °C overnight, then oocytes were washed by wash buffer (0.1% Tween 20 and 0.01% Triton X-100 in PBS) for 3 times (5 min each time). Next the oocytes were labeled with secondary antibody coupled to FITC-conjugated goat anti-rabbit IgG (1:100) at room temperature for 1 h. For α-tubulin staining, MI stage oocytes were incubated with anti-α-tubulin-FITC antibody (1:200). For actin staining, GV and MI stage oocytes were incubated with Phalloidin-TRITC at room temperature for 2 h. Then the oocytes were washed as the same way. Finally, oocytes were incubated with Hoechst 33342 at room temperature for 10-20 min. After staining, samples were mounted on glass slides and observed with a confocal laser-scanning microscope (Zeiss LSM 800 META, Germany).

### ER and Mito-tracker staining

To study ER and mitochondria distribution during mouse oocyte meiosis, MI stage oocytes were incubated with ER-Tracker Red (1:3000) or 200 nM Mito-tracker green (Red) in M16 medium for 30 min at 37°C and 5% CO2. Then the oocytes were washed three times with M2 medium, finally the samples were examined with confocal microscopy. During the MI stage of oocyte development, both the ER and mitochondria evenly distributed in the cytoplasm and accumulated at the spindle periphery in MI stage. This clustering phenomenon is considered as the normal localization pattern of these organelles. On the other hand, when the ER and mitochondria cluster randomly within the cytoplasmic region, this localization is defined as organelle abnormal localization pattern.

### JC-1 detection

The enhanced mitochondrial membrane potential assay Kit was employed to analyze the mitochondrial membrane potential of oocytes. The MI stage oocytes were transferred from the M16 medium to JC-1 for 30 min at 37°C and 5% CO2. Following three washes with M2 medium, the oocytes were examined using a fluorescent microscope (OLYMPUS IX71, Japan) for the presence of a fluorescent signal.

### Time lapse microscopy

To image the dynamic changes that occurred during oocyte maturation, oocytes were cultured in M16 medium, then transferred to the Leica SD AF confocal imaging system equipped with 37 °C incubator and 5% CO2 supply (H301-K-FRAME). The spindle in oocytes was labeled by α-tubulin-EGFP.

### Immunoprecipitation

4-6 ovaries were put into RIPA Lysis Buffer contained phosphatase inhibitor cocktail (100×) (Kangwei Biotechnology, China), and were completely cleaved on ice block. We collected supernatant after centrifugation (13200 rps, 20 min) and then took out 50 μl as input sample at 4 °C. The rest of the supernatant was incubated with primary antibody (FMNL2 or INF2 antibody) overnight at 4 °C. 30 μl conjugated beads (washed five times in PBS) were added to the supernatant/antibody mixture and incubated at 4 °C for 4-6 h, after three times wash by immune complexes, the samples were then released from the beads by mixing in 2× SDS loading buffer for 10min at 30 °C.

### Western blot analysis

Approximate 100-150 GV or MI stage mouse oocytes were placed in Laemmli sample buffer and heated at 85l for 7-10 min. Proteins were separated by SDS-PAGE at 165V for 70-80 min and then electrophoretically transferred to polyvinylidene fluoride (PVDF) membranes (Millipore, Billerica, MA, USA) at 20 V for 1 hour. After transfer, the membranes were then blocked with TBST (TBS containing 0.1% Tween 20) containing 5% non-fat milk at room temperature for 90 min. After blocking, the membranes were incubated with rabbit monoclonal anti-FMNL2 antibody (1:500), rabbit monoclonal anti-Arp2 antibody (1:500), mouse monoclonal anti-profilin1 antibody (1:500), rabbit monoclonal anti-Fascin antibody (1:5000), rabbit polyclonal anti-INF2 antibody (1:500), rabbit monoclonal anti-Grp78 antibody (1:1000), rabbit monoclonal anti-cofilin antibody (1:2000), rabbit monoclonal anti-CHOP antibody (1:1000), or rabbit monoclonal anti-tubulin antibody (1:2000) at 4 °C overnight. After washing 5 times in TBST (5 min each), membranes were incubated for 1h at room temperature with HRP-conjugated Pierce Goat anti-Rabbit IgG (1:5000) or HRP-conjugated Pierce Goat anti-mouse IgG (1:5000). After washing for 5 times, the membranes were visualized using chemiluminescence reagent (Millipore, Billerica, MA). Every experiment repeated at least 3 times with different samples.

### Fluorescence Intensity Analysis

Immunofluorescence experiments were conducted simultaneously and with consistent parameters in both the control and treatment groups. The images were consistently captured using identical confocal microscope settings. Subsequently, the average intensity of fluorescence per unit area in the designated region of interest was quantified following the fluorescence staining. The acquired fluorescence data was analyzed employing ZEN 2011 and ImageJ software.

### Statistical analysis

All statistical analyses were performed using GraphPad Prism7.00 software (GraphPad, CA, USA), employing the t-test to assess the statistical significance between the control and treatment groups. The results were represented as the mean ± standard error of the mean (SEM). Statistical significance was defined as a P-value < 0.05, denoted as *, ** for P < 0.01, ***, and **** for P < 0.001 and P < 0.0001, respectively. Every experiment was conducted with a minimum of three biological replicates.

## Results

### Expression and subcellular localization of FMNL2 during mouse oocyte maturation

We first examined FMNL2 expression in mouse oocytes at different stages. The results indicated that FMNL2 all expressed in GV, MI and MII stages during mouse oocyte maturation (GV, 1; MI, 0.82 ± 0.07; MII, 0.61 ± 0.10, Figure 1A). Next, we performed FMNL2-EGFP mRNA microinjection to examine the localization of FMNL2. As shown in Figure 1B, FMNL2 accumulated at the oocyte cortex during the GV, GVBD MI and MII stages. Besides, FMNL2 also localized at the spindle periphery during MI stages. The FMNL2 antibody staining results also confirmed this localization pattern. In addition, we co-stained FMNL2 antibody with F-actin, and the results revealed that both FMNL2 and F-actin are localized in the cortex region of oocytes (Figure 1C). The FMNL2 localization pattern indicated that FMNL2 might interact with actin dynamics during oocyte meiosis.

**Figure 1.**
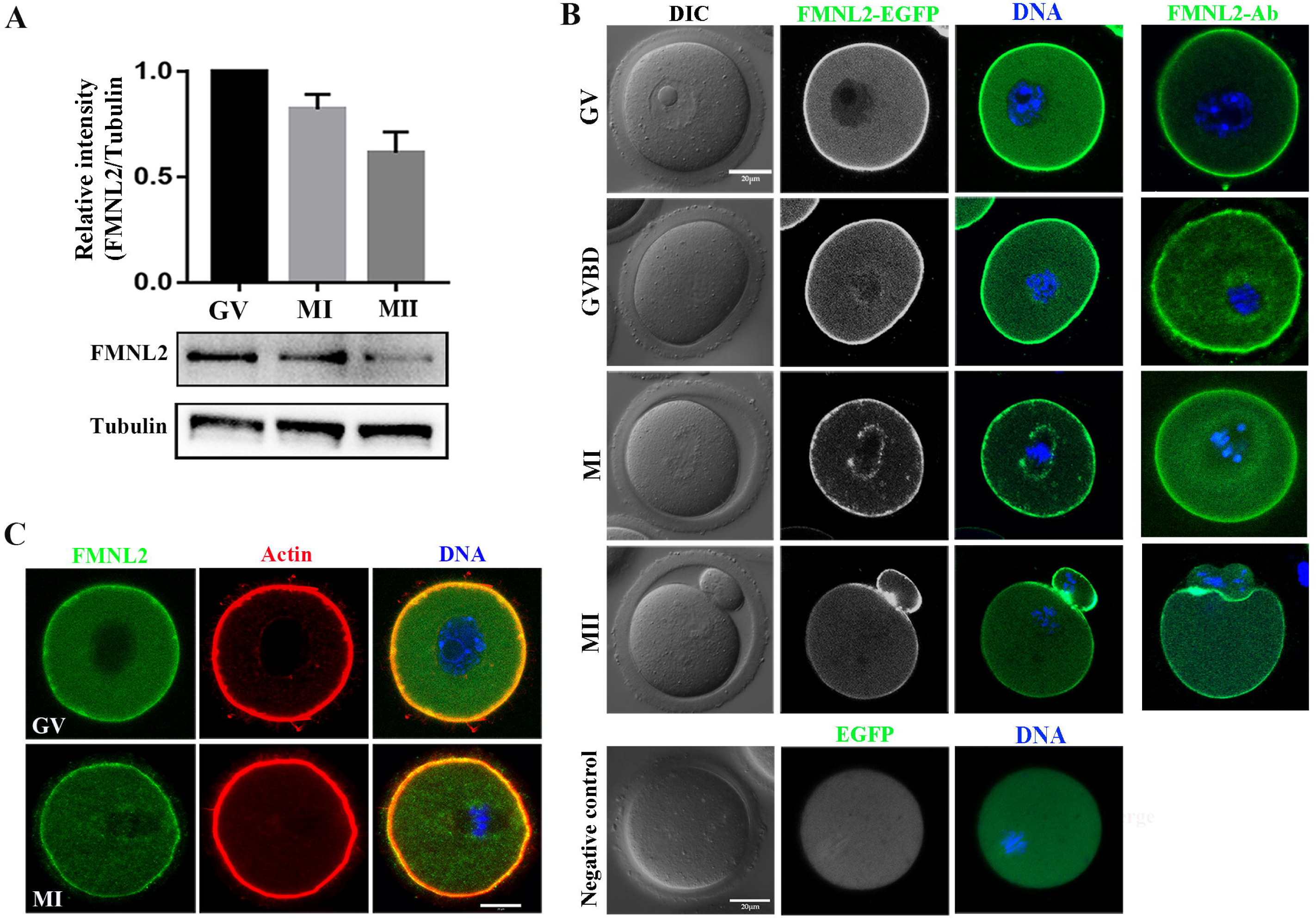
Expression and subcellular localization of FMNL2 during mouse oocyte meiosis. **(A)** Western blotting results of FMNL2 protein expression at different stages. FMNL2 expressed at the GV, MI, and MII stages. **(B)** Subcellular localization of FMNL2-EGFP and FMNL2 antibody during mouse oocyte meiosis. FMNL2 was enriched at the cortex (GV, GVBD, MI and MII stage) and spindle periphery (MI stage). Green, FMNL2-EGFP; blue, DNA. Negative control: Green, EGFP; blue, DNA. Bar =20 μm. **(C)** Co-staining of oocytes for FMNL2 and actin. FMNL2 and actin both localization in cortex. Green, FMNL2-antibody; red, actin; blue, DNA. Bar =20 μm.

### FMNL2 is essential for polar body extrusion and asymmetric division in mouse oocytes

To investigate the functional roles of FMNL2 in mouse oocytes, we employed FMNL2 siRNA microinjection to knockdown FMNL2 protein expression. A significant decrease of FMNL2 protein level was shown in FMNL2-KD oocytes compared to control group by western blot (1 vs. 0.48 ± 0.08, P < 0.01, Figure 2A). We then examined the maturation of oocytes, and the results indicated that knockdown of FMNL2 had an impact on the extrusion of the first polar body. Moreover, a significant proportion of oocytes exhibited larger polar bodies upon extrusion (Figure 2B). Based on the size of the extruded polar bodies, those with a diameter exceeding one-third of the oocyte’s diameter were categorized as large polar bodies. Consequently, we proceeded to calculate the rates of polar body extrusion and the generation of large polar bodies in the oocytes. The quantitative results also confirmed this phenotype (rate of polar body extrusion: 74.26 ± 1.44%, n = 439 vs. 59.5 ± 2.82%, n = 398, P < 0.001, Figure 2C; rate of large polar bodies: 19.05 ± 1.97%, n = 311 vs. 37.16 ± 1.87%, n = 257, P < 0.0001, Figure 2D). In addition, live-cell imaging was used to determine the dynamic changes that occurred during oocyte maturation, and the results showed that the oocytes either failed to undergo cytokinesis or divided from the central axis of the oocytes and formed big polar bodies (Figure 2E). To further confirm the phenotype, we performed FMNL2 rescue experiments by expressing exogenous Fmnl2 mRNA in FMNL2-depleted oocytes (Figure 2F), we found that exogenous Fmnl2 mRNA expression rescued first polar body extrusion and large polar body defects (Figure 2G). The quantitative results also confirmed this phenotype (rate of polar body extrusion: 48.34 ± 4.2%, n = 355 vs. 62.62 ± 3.6%, n = 377, P < 0.01, Figure 2H; rate of large polar bodies: 30.93 ± 2%, n = 193 vs. 9.58 ± 2.4%, n = 203, P < 0.01, Figure 2I). It is known that knockdown of FMNL3 leads to inhibition of oocyte maturation. To investigate whether FMNL2 exhibits an additive effect with FMNL3 in terms of functionality, we simultaneously knocked down both FMNL2 and FMNL3. The results demonstrated that simultaneous knockdown of the two FMNL proteins, compared to the control group, resulted in a decrease in oocyte maturation rate. However, when compared to the single knockdown of FMNL2, the double knockdown of FMNL2 and FMNL3 did not cause more severe defects in polar body extrusion. (polar body extrusion, Control: 70.97 ± 1.23%, n=261 vs FMNL2+3-KD: 60.42 ± 2.99%, n=198, P < 0.05, Figure 2J. Large polar body, Control: 10.85 ± 0.97%, n=172 vs FMNL2+3-KD: 32.90 ± 1.88%, n=118, P < 0.001, Figure 2K). These results suggested that FMNL2 played critical roles for the polar body extrusion and asymmetric division during mouse oocyte maturation.

**Figure 2.**
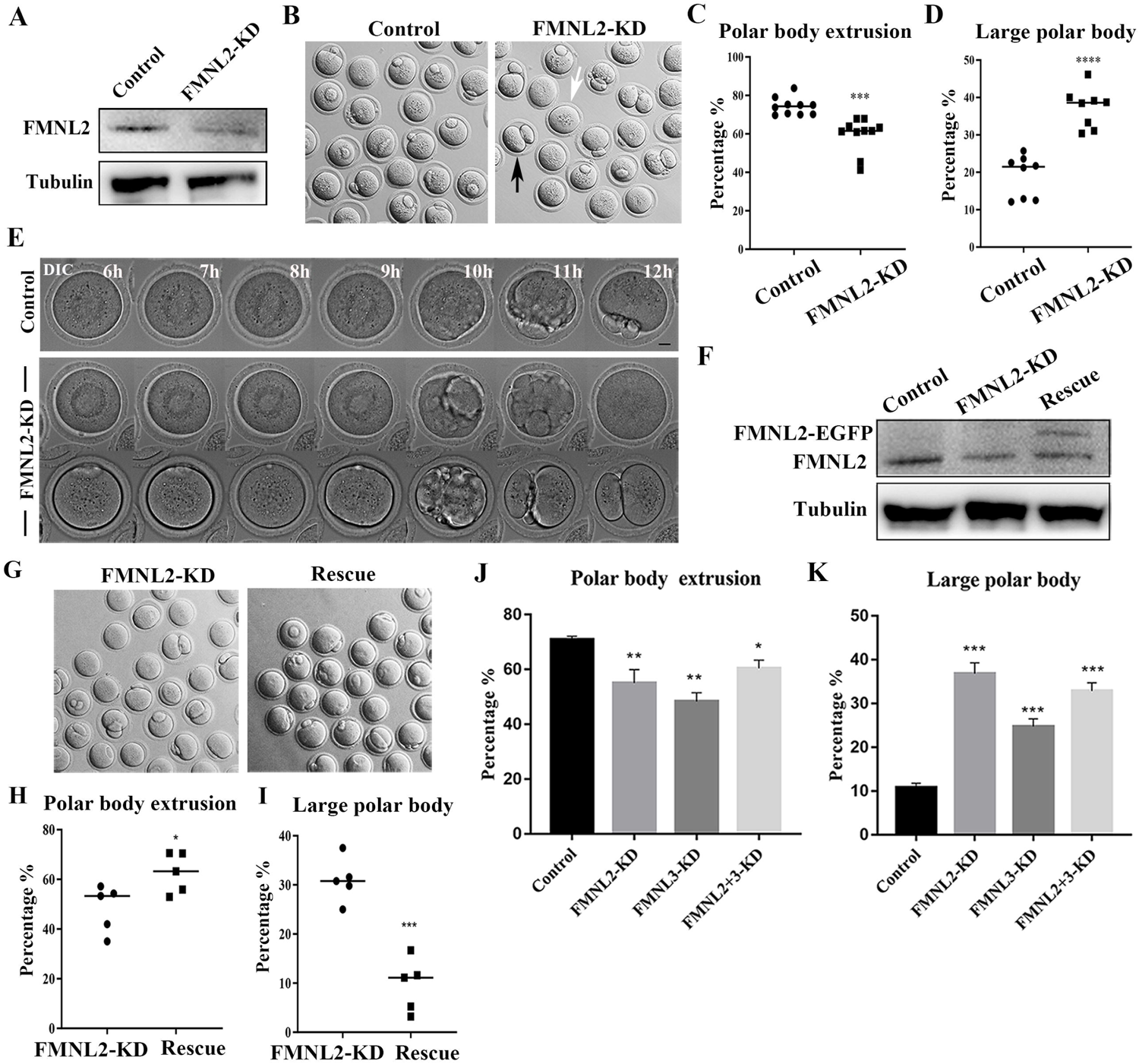
Knockdown of FMNL2 affects first polar body extrusion and asymmetric division. **(A)** Western blot analysis for FMNL2 expression in the FMNL2-KD group and control group. Relative intensity of FMNL2 and tubulin was assessed by densitometry. **(B)** DIC images of control oocytes and FMNL2-KD oocytes after 12 h culture. FMNL2-KD caused large polar bodies (black arrows) and some oocytes failed to extrude the polar bodies (white arrows). (**C)** Rate of polar body extrusion after 12 h culture of the control group and FMNL2-KD group. **(D)** Rate of large polar body extrusion after 12 h culture in the control group and FMNL2-KD group. **(E)** Time-lapse microscopy showed that polar body extrusion failed after FMNL2-KD. Bar = 10 μm. **(F)** Western blot analysis for FMNL2 expression in the control group, FMNL2-KD group and rescue group. Relative intensity of FMNL2 and tubulin was assessed by densitometry. **(G)** DIC images of FMNL2-KD oocytes and rescue oocytes after 12 h culture. (**H)** Rate of polar body extrusion after 12 h culture of the FMNL2-KD group and rescue group. **(I)** Rate of large polar body extrusion after 12 h culture in the FMNL2-KD group and rescue group. (**J)** Rate of polar body extrusion after 12 h culture of the control group and FMNL2+3-KD group. **(K)** Rate of large polar body extrusion after 12 h culture in the control group and FMNL2+3-KD group. The data are presented as mean ± SEM from at least three independent experiments. * P < 0.05, ** P < 0.01, *** P < 0.001, **** P < 0.0001.

### FMNL2 regulates spindle migration during mouse oocyte maturation

To investigate the causes for polar body extrusion defects, we examined the spindle migration by time-lapse microscopy during oocyte meiosis. As shown in Figure 3A, in the control oocyte, the meiotic spindle formed in the center of the oocyte after culture 8 h and moved to the oocyte cortex at 9.5h; and the polar body was extruded at 11-12h, with a spindle formed near the cortex at MII stage. However, in FMNL2-KD oocytes, two phenotypes were observed: 1) the meiotic spindle remained in the center of the oocyte until 10 h, and then the oocytes initiated the cytokinesis at 10.5 h but failed to extrude the polar body; 2) some oocytes with arrested spindles initiated the cytokinesis but extruded a big polar body (Figure 3A). This indicated the failure of spindle migration after FMNL2 depletion. We analyzed the rate of cortex-localized spindle in oocytes by cultured for 9.5 h, and the result showed that the rate of migrated spindles in control oocytes was significantly higher than that in FMNL2-KD oocytes (59.94 ± 3.42%, n = 78 vs. 38.97 ± 6.34%, n = 64, P < 0.05, Figure 3B). We also performed FMNL2 rescue experiments. Supplementing with exogenous Fmnl2 rescued the spindle migration defects compared with the FMNL2-depletion group (40.27 ± 3.19%, n = 181 vs. 57.01± 2.72%, n = 57, P < 0.01, Figure 3C). These results suggested that FMNL2 might be involved in spindle migration in mouse oocytes.

**Figure 3.**
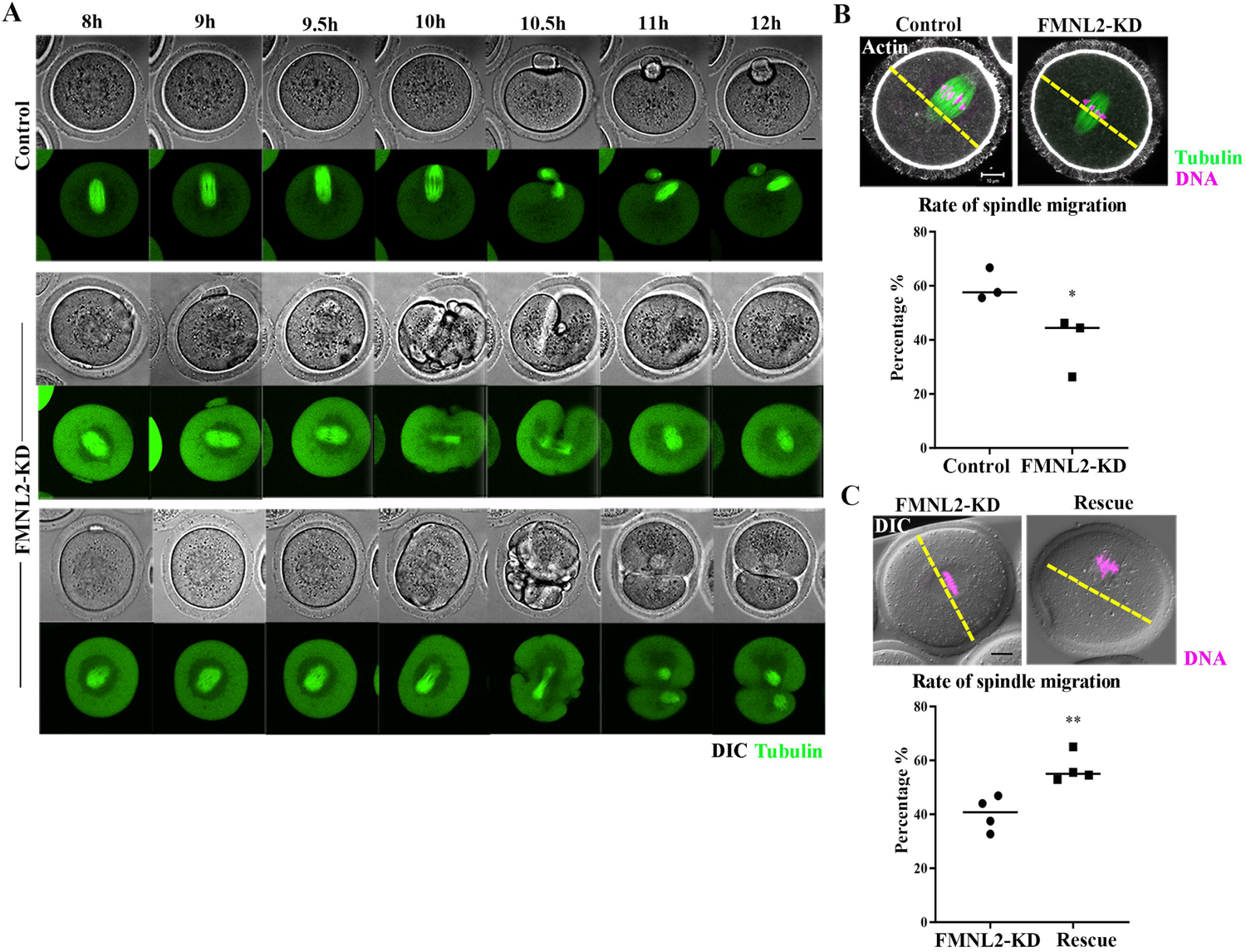
Knockdown of FMNL2 disrupts spindle localization during mouse oocyte meiosis. **(A)** Time-lapse microscopy showed that spindle migration failed after FMNL2-KD. Green, tubulin-EGFP. Bar = 10 μm. **(B)** Representative images and the proportion of spindle migration after 9.5 h of culture in the control group and FMNL2-KD oocyte group. White, actin; green, tubulin; magenta, DNA. Bar = 10 μm. **(C)** Representative images and the proportion of spindle migration after 9.5 h of culture in the FMNL2-KD group and rescue oocyte group. magenta, DNA. Bar = 10 μm. The data are presented as mean ± SEM from at least three independent experiments. * P < 0.05, ** P < 0.01.

### FMNL2 promotes cytoplasmic actin assembly during mouse oocyte maturation

As FMNL2 is a key actin assembly factor, we further investigated actin assembly after deleting FMNL2 in mouse oocytes. Surprisingly there was no significant difference for the signals of cortex actin was observed between control oocytes and FMNL2-KD oocytes, which was confirmed by the fluorescence intensity analysis (30.88 ± 1.10, n = 28 vs. 30.58 ± 1.12, n = 28, P > 0.05, Figure 4A, 4B). However, we found a significant decrease of cytoplasmic actin signals in the FMNL2-KD oocytes, and the statistical analysis for the cytoplasmic actin fluorescent signals also confirmed our findings (58.25 ± 2.05, n = 26 vs. 37.92 ± 2.02, n = 24, P < 0.0001, Figure 4C, 4D). Moreover, the rescue experiments showed that exogenous FMNL2 rescued the decrease of cytoplasmic actin filaments compared with the FMNL2-depletion group (37.98 ± 1.98, n = 16 vs. 54.72± 2.88, n = 15, P < 0.0001, Figure 4E, 4F). We next explored how FMNL2 regulates cytoplasmic actin assembly in oocytes. By mass spectrometry analysis we found there were several actin-related potential candidates which might be related with FMNL2 (Figure 4G). Co-immunoprecipitation results showed that FMNL2 precipitated Arp2 and Formin2 but not Profilin and fascin (Figure 4H). To further verify the correlation between FMNL2 and Arp2 and Formin2, we then examined Arp2 and Formin2 protein expression after FMNL2 knockdown. The results showed Arp2 protein expression increased significantly after FMNL2 knockdown (1 vs. 1.56 ± 0.07, P < 0.001, Figure 4I) but Formin2 decreased after FMNL2 knockdown (1 vs. 0.62 ± 0.04, P < 0.001, Figure 4J). Exogenous FMNL2 rescued these alterations compared with that in the FMNL2-KD group (Arp2 protein expression: 1 vs. 0.65 ± 0.06, P < 0.01, Figure 4I; Formin2 protein expression: 1 vs. 1.24 ± 0.05, P < 0.01, Figure 4J). These results indicated that FMNL2 may be associated with Formin2 and Arp2 for actin assembly in mouse oocytes.

**Figure 4.**
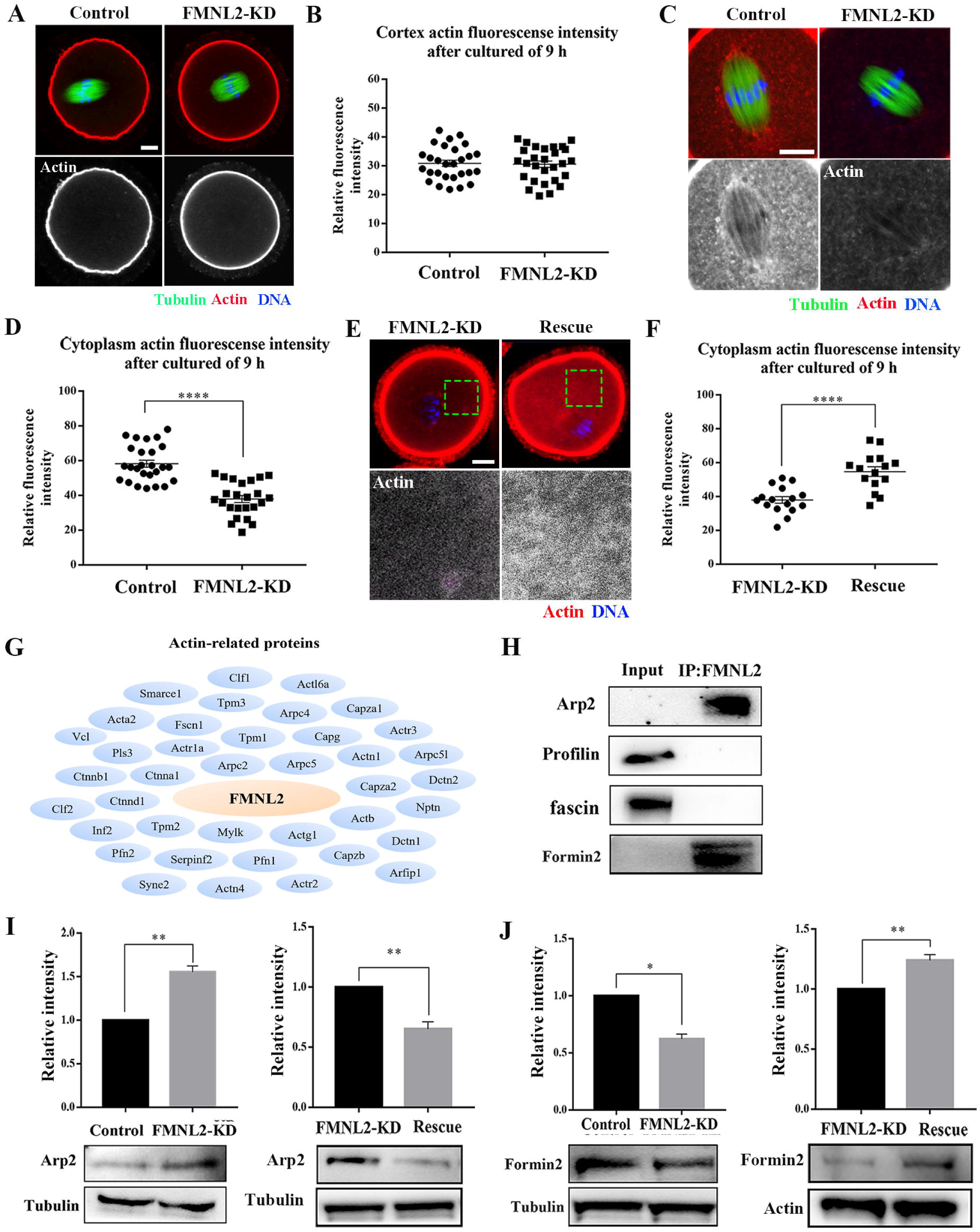
Knockdown of FMNL2 disrupts actin assembly during mouse oocyte meiosis. **(A, B)** Representative images of actin distribution at the oocyte cortex and the fluorescent intensities in the control group and FMNL2-KD group (P > 0.1). White, actin; green, tubulin; magenta, DNA. Bar = 10 μm. **(C, D)** Representative images of actin distribution in the oocyte cytoplasm and the fluorescent intensities in the control group and FMNL2-KD group. White, actin; green, tubulin; magenta, DNA. Bar = 10 μm. **(E, F)** Representative images of actin distribution in the oocyte cytoplasm and the fluorescent intensities in the FMNL2-KD group and rescue group. White, actin; magenta, DNA. Bar = 10 μm. **(G)** Mass spectrometry results showed that FMNL2 was related to many actin-related proteins. **(H)** Co-IP results showed that FMNL2 was correlated with Arp and Formin2 but not with Profiling and Fascin. **(I)** Arp2 protein expression significantly increased in the FMNL2-KD oocytes compared with the control oocytes. Arp2 protein expression significantly decreased in the rescue oocytes compared with the FMNL2-KD oocytes. **(J)** Formin2 protein expression significantly decreased in the FMNL2-KD oocytes compared with the control oocytes. Formin2 protein expression significantly increased in the rescue oocytes compared with the FMNL2-KD oocytes. The data are presented as mean ± SEM from at least three independent experiments. * P < 0.05, ** P < 0.01, **** P < 0.0001.

### FMNL2 regulates endoplasmic reticulum distribution during mouse oocyte maturation

The mass spectrometry analysis data indicated that several ER-related potential candidates which might be related with FMNL2 (Figure 5A), while INF2, a typical protein which mediates actin polymerization at ER showed high confidence level. We then examined the relationship between FMNL2 and INF2, and the co-immunoprecipitation results showed that FMNL2 precipitated INF2 and INF2 also precipitated FMNL2 (Figure 5B), indicating that FMNL2 interacted with INF2 in mouse oocytes. We then examined the ER distribution in FMNL2-KD oocytes. As shown in Figure 5C, in control oocytes the ER evenly distributed in the cytoplasm and accumulated at the spindle periphery in MI stage; however, ER agglomerated in cytoplasm in FMNL2-KD oocytes (Figure 5C). The statistical analysis showed that the abnormal distribution of ER increased significantly in the FMNL2-KD group (28.91 ± 5.62, n = 27 vs. 59.64 ± 6.95, n = 28, P < 0.05, Figure 5D). The localization pattern of ER indicated its functions might be disturbed. In FMNL2-KD oocytes, we found the expressions of ER-stress related proteins Grp78 and Chop were significantly increased (Grp78: 1 vs. 1.42 ± 0.12, P < 0.05; Chop: 1 vs. 1.53 ± 0.16, P < 0.05. Figure 5E), indicating the occurrence of ER stress. We also performed FMNL2 rescue experiments. Supplementing with exogenous Fmnl2 rescued the ER distribution defects caused by FMNL2 knockdown (Figure 5F), which was supported by the statistical analysis showing that the abnormal distribution rate of ER decreased significantly in the rescue group (52.04 ± 5.29, n = 70 vs. 34.91 ± 3.37, n = 78, P < 0.05, Figure 5G). Moreover, Grp78 protein expression decreased in the rescue group (1 vs. 0.78 ± 0.05, P < 0.01. Figure 5H). These results indicated that the depletion of FMNL2 affected ER distribution and caused ER stress in mouse oocytes.

**Figure 5.**
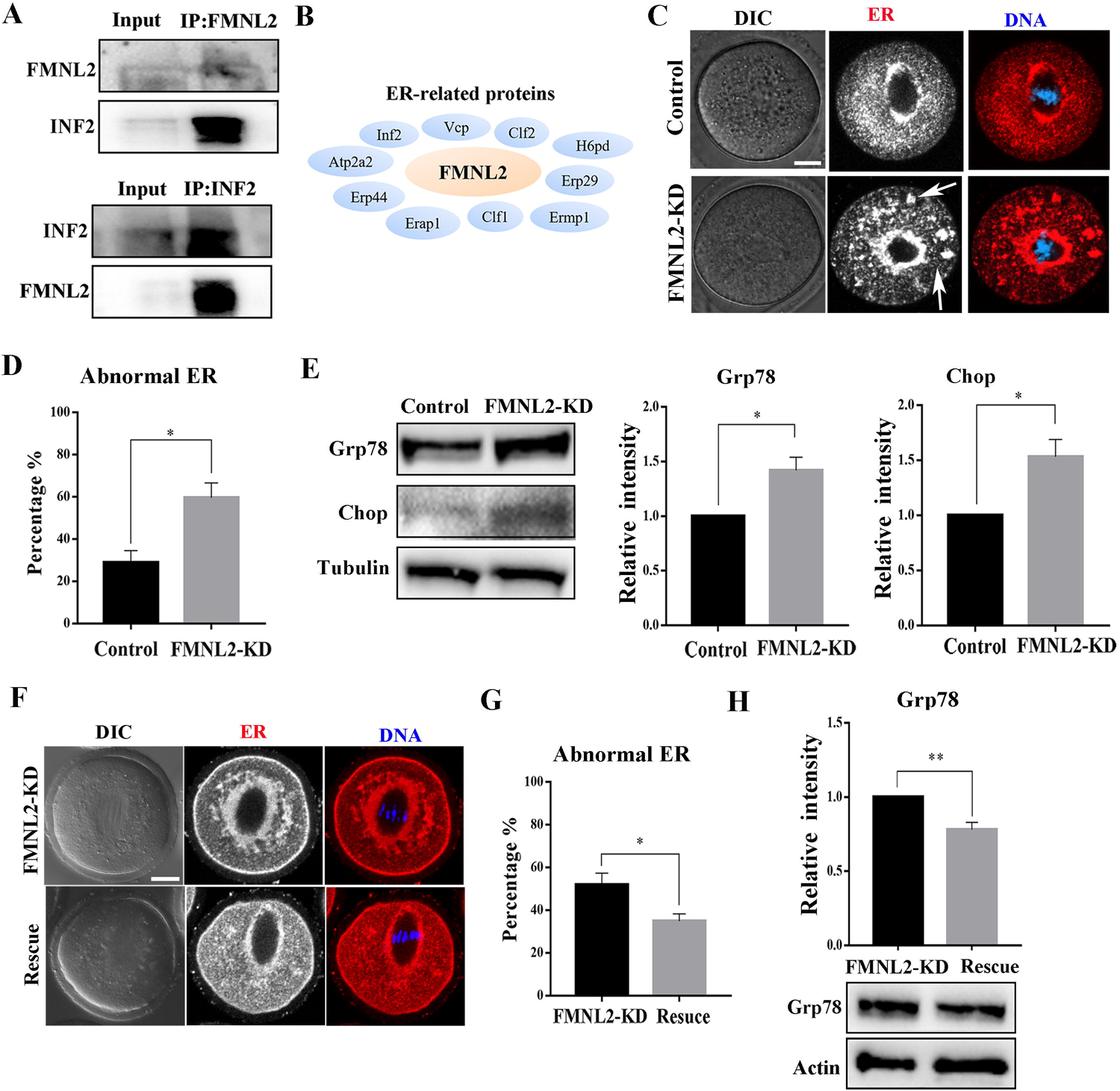
FMNL2 regulates endoplasmic reticulum distribution during mouse oocytes maturation. **(A)** Co-IP results showed that FMNL2 was correlated with INF2. **(B)** Mass spectrometry results showed that FMNL2 was associated with ER-related proteins. **(C)** Representative images of ER distribution in the oocyte cytoplasm in the control group and FMNL2-KD group. In FMNL2-KD oocytes, ER agglomerated in cytoplasm (white arrow). Red, ER; Blue, DNA. Bar = 10 μm. **(D)** Abnormal distribution of ER significantly increased in the FMNL2-KD oocytes compared with the control oocytes. **(E)** Grp78 and Chop protein expression significantly increased in the FMNL2-KD oocytes compared with the control oocytes. The band intensity analysis also confirmed this finding. **(F)** Representative images of ER distribution in the oocyte cytoplasm in the FMNL2-KD group and rescue group. Red, ER; Blue, DNA. Bar = 10 μm. **(G)** Abnormal distribution of ER significantly decreased in the rescue oocytes compared with the FMNL2-KD oocytes. **(H)** Grp78 protein expression significantly decreased in the rescue oocytes compared with the FMNL2-KD oocytes. The band intensity analysis also confirmed this finding. The data are presented as mean ± SEM from at least three independent experiments. * P < 0.05, ** P < 0.01.

### FMNL2 regulates mitochondrial distribution during mouse oocyte maturation

As INF2 is also related to the mitochondrial connection of ER, we further screened up the mass spectrometry analysis data and we found many mitochondria-related potential candidates which might be related with FMNL2 (Figure 6A). Therefore, we further examined the distribution of mitochondria in FMNL2-KD oocytes. In control oocytes, the mitochondria evenly distributed in the cytoplasm and accumulated at the spindle periphery in MI stage; however, in FMNL2-KD oocytes, mitochondria presented clumped aggregation distribution in cytoplasm (Figure 6B). We counted the number of clumps and found that the uniform distribution of mitochondria decreased significantly in the FMNL2-KD group (59.66 ± 8.48, n = 31 vs. 20.83 ± 4.17, n = 32, P < 0.05, Figure 6C). A large number of FMNL2-KD oocytes agglomerated into one to three clumps (22.73± 4.27, n = 31 vs. 42.50 ± 1.25, n = 32, P < 0.05, Figure 6C). Supplementing with exogenous Fmnl2 rescued the mitochondria distribution (Figure 6D), the statistical analysis showed that the uniform distribution of mitochondria increased significantly in the rescue group (36.49 ± 3.97, n = 53 vs. 53.90 ± 2.09, n = 79, P < 0.05, Figure 6E). We also examined mitochondrial membrane potential, and the results showed that FMNL2 depletion caused the alterations of mitochondrial membrane potential (MMP) by JC-1 staining. The fluorescence intensity of JC-1 red channel was decreased compared with the control group (Figure 6F). We also calculated the ratio for red/green fluorescence intensity, and the results also confirmed this (control group: 0.40 vs. FMNL2-KD: 0.21 ± 0.01, P < 0.01) (Figure 6G). Cofilin is an important factor of actin assembly and regulates mitochondrial function. We also examined cofilin protein expression after FMNL2 knockdown. The results showed cofilin protein expression decreased significantly after FMNL2 knockdown (1 vs. 0.81 ± 0.03, P < 0.01, Figure 6H). These results indicated that FMNL2 regulated mitochondria distribution and function during mouse oocyte maturation.

**Figure 6.**
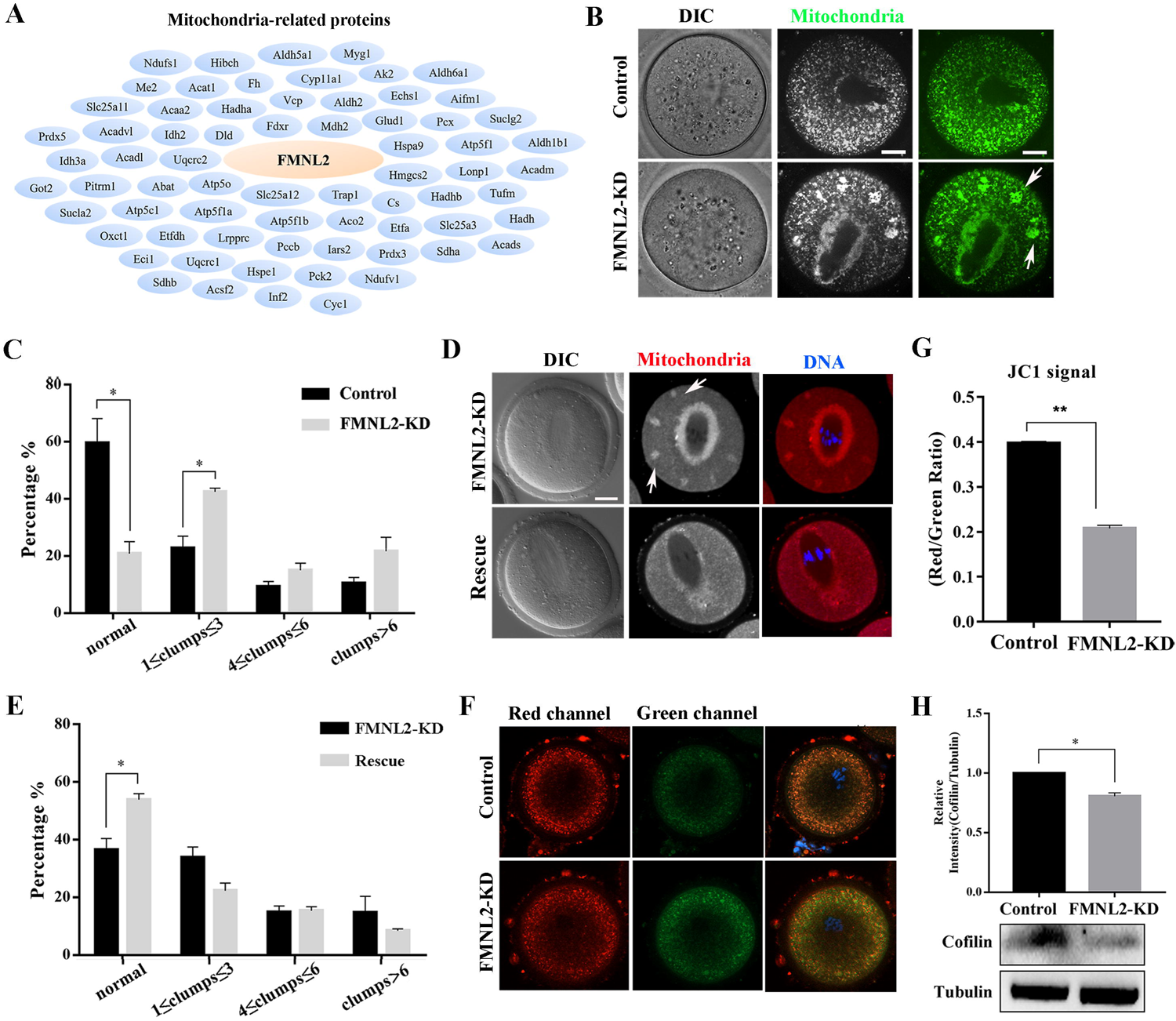
FMNL2 regulates mitochondrial distribution during mouse oocytes maturation. **(A)** Mass spectrometry results showed that FMNL2 was related to many mitochondria-related proteins. **(B)** Representative images of mitochondrial distribution in the oocyte cytoplasm in the control group and FMNL2-KD group. In FMNL2-KD oocytes, mitochondrial agglomerated in cytoplasm (white arrow). Green, Mito. Bar = 20 μm. **(C)** Abnormal distribution of mitochondrial significantly increased in the FMNL2-KD oocytes compared with the control oocytes. **(D)** Representative images of mitochondrial distribution in the oocyte cytoplasm in the FMNL2-KD group and rescue group. In FMNL2-KD oocytes, mitochondrial agglomerated in cytoplasm (white arrow). Red, Mito; Blue, DNA. Bar = 20 μm. **(E)** Abnormal distribution of mitochondrial significantly decreased in the rescue oocytes compared with the FMNL2-KD oocytes. **(F)** The typical picture for JC1 green channel and red channel after FMNL2-KD. **(G)** The JC1 signal (red/green ratio) after FMNL2-KD compare with the control group, the JC-1 red/green fluorescence ratio was significantly reduced in FMNL2-KD groups. Bar = 20 µm. **(H)** cofilin protein expression significantly decreased in the FMNL2-KD oocytes compared with the control oocytes. The band intensity analysis also confirmed this finding. The data are presented as mean ± SEM from at least three independent experiments. * P < 0.05, ** P < 0.01.

## Discussion

In this study, we explored the functions of FMNL2 during mouse oocyte meiosis. Our results indicated that FMNL2 regulated actin-based spindle migration for asymmetric cell division of oocytes, and more importantly FMNL2 was critical for maintaining the distribution of the ER and mitochondria, which set up a link for actin-related spindle migration and organelle dynamics in oocytes.

As a subfamily of Formin family, FMNLs play an important role in regulating actin filaments (*18*), while FMNL2 is most widely expressed in variety of cell models among the members of FMNLs. In this study, we showed that FMNL2 expressed in mouse oocytes and it mainly accumulated at the oocyte cortex and spindle periphery, which was similar with the actin distribution pattern in oocytes. This specific localization is also similar to FMN2, a well-studied factor in the formin family for spindle migration during oocyte meiosis (*7, 19*). In addition, another FMNLs family member, FMNL1 is also localized at the cortex and is essential for actin polymerization and spindle assembly during oocyte meiosis (*20*). Based on the localization pattern of FMNL2, we speculated that the functions of FMNL2 might be also involved in actin-related process during mouse oocyte meiosis.

To confirm our hypothesis, we depleted FMNL2 protein expression and we found that absence of FMNL2 caused the aberrant first polar body extrusion. The oocytes either failed to form the polar body or extruded large polar bodies. These phenotypes caused by FMNL2 depletion are similar to the other actin-related proteins during oocyte maturation such as Arp2/3 complex (*8, 21*) and FMN2 (*22, 23*). We next examined the actin distribution in oocytes since it is reported that FMNL2 promotes actin filament assembly in many models. FMNL2 is required for cell-cell adhesion formation by regulating the actin assembly (*24*), and FMNL2 could directly drives actin elongation (*15*). In CRC cells, cortactin bind to FMNL2 to active the actin polymerization, and FMNL2 is important for invadopodia formation and functions (*25*). Our results showed that the FMNL2 depletion caused significantly decrease in cytoplasmic actin, indicating the conserved roles of FMNL2 on actin assembly in mammalian oocyte model. Other Formin family proteins such as Daam1, FHOD1, and Formin-homology family protein mDia1 are also reported to affect oocyte meiosis by regulating actin polymerization (*26–28*).

We then tried to explore how FMNL2 involves into the actin assembly in oocytes. Mass spectrometry analysis data indicated that FMNL2 associated with several actin-related proteins, and we found that a potential association between FMNL2 and Arp2/Formin2. This could be confirmed by the altered expression of these two molecules after FMNL2 depletion. Therefore, we speculated FMNL2 could regulate cytoplasmic actin assembly in oocytes through the association with Formin2 since it is reported to be an important protein for cytoplasmic actin assembly in oocytes (*22*). Interestingly, our results showed that unlike the reduction of cytoplasmic actin, cortex actin was not affected by the absence of FMNL2. We speculate that FMNL2 and Arp2/3 both contribute to the cortex actin dynamics, when FMNL2 decreases, ARP2 increases to compensate for this, which maintains the cortex actin level. As an actin nucleator Arp2/3 complex localizes at the cortex and is essential for actin polymerization during oocyte meiosis (*8, 29*). These results suggested that FMNL2 might be involved in cytokinesis and asymmetric division by regulating actin assembly during mouse oocyte maturation.

The spindle migration is a key step in ensuring the asymmetric division for oocytes (*30*). In mitosis, spindle position is decided by cortical actin and astral microtubules; in contrast, spindle migration is mainly mediated by actin filaments during oocyte meiosis (*30, 31*). Due to the effects of FMNL2 on asymmetric division and cytoplasmic actin, we analyzed the spindle positioning at late MI, we found that the spindle migration was disturbed after FMNL2 depletion, no matter the cytokinesis occurred or not. Several formin proteins are shown to regulate spindle migration during oocytes meiosis. For example, FMN2 nucleates actin surrounding the spindle, pushing force generated by actin to trigger the spindle migration (*19, 22*), and cyclin-dependent kinase 1 (Cdk1) induces cytoplasmic Formin-mediated F-actin polymerization to propel the spindle into the cortex (*32*). Our previous studies also showed that absence of the formin family member FMNL1 or FHOD1 could lead to the decrease of cytoplasmic actin to prevent the spindle migration (*20, 27*). We speculated that FMNL2 together with other Formin proteins, conservatively regulate actin-mediated spindle migration during oocyte meiosis.

Another important finding is that through the mass spectrometry analysis we found many candidate proteins which were related with ER, and our results indicated that FMNL2 was essential for the maintenance of ER distribution in the cytoplasm. Moreover, the loss of FMNL2 induced ER stress, showing with altered expression of GRP78 and CHOP. Proper distribution of ER is important for the oocyte quality. ER displays a homogeneous distribution pattern throughout the entire ooplasm during development of oocytes and embryos from diabetic mice(*33*). During the transition of mouse oocytes from MI to MII phase, actin regulates cortical ER aggregation(*34*). In addition, Formin2 is shown to colocalize with the ER during oocyte meiosis and the ER-associated Formin2 at the spindle periphery is required for MI chromosome migration(*6*). In our results we showed that FMNL2 associated with INF2 protein in oocytes. INF2 is an ER-associated protein, and the expression of GFP-INF2 which containing DAD/WH2 mutations causes the ER to collapse around the nucleus (*35*). We concluded that FMNL2 might regulate INF2 for the distribution of ER in cytoplasm of oocytes.

Besides its roles of ER distribution, it is shown that INF2 also affects mitochondrial length and ER-mitochondrial interaction in an actin-dependent manner (*35, 36*). It is shown that INF2 regulates Drp1 for mitochondrial fission, and INF2-induced actin filaments may drive initial mitochondrial constriction, which allows Drp1-driven secondary constriction(*36, 37*). In addition, we also found many candidate proteins which were related with mitochondria from mass spectrometry analysis. During oocyte meiosis, mitochondria gradually accumulated around the spindle after GVBD, and the spindle-peripheral FMN2 and its actin nucleation activity are important for the accumulation of mitochondria in this region (*19*). Our results found that FMNL2 depletion caused agglutination of mitochondria and altered MMP level in the cytoplasm, indicating its roles on the mitochondria distribution and functions. Another formin protein mDia1 is shown to be necessary to induce the anchoring of mitochondria along the cytoskeletal in mammalian CV-1 cells and Drosophila BG2-C2 neuronal cells(*38*). Moreover, the formin interaction protein Spire1C binds INF2 to promote actin assembly on mitochondrial surfaces, and Spire1C disruption could reduce mitochondrial constriction and division(*39*). In addition, our result indicated that cofilin expression decreased in FMNL2 depletion oocytes. Cofilin is an actin-depolymerizing factor and its localization at the mitochondrial fission site is crucial for inducing mitochondrial fission and mitophagy (*40*). Depleting of cofilin resulted in abnormal interconnection and elongation of mitochondria (*41*). Together with its roles on ER, these data indicated that FMNL2 might associate with INF2 and cofilin for the actin-based organelle distribution during oocyte meiosis.

Collectively, we provide a body of evidence showing that FMNL2 associates with Formin2 and Arp2/3 complex for actin assembly, which further regulates spindle migration and INF2/Cofilin-related organelle dynamics during mouse oocyte maturation.

## Data Availability

All data generated or analyzed during this study are included in this published article

## Acknowledgement

We are particularly grateful to Xiao-Yan Fan and Xing-Hua Wang from Fertility Preservation Laboratory, Reproductive Medicine Center, Guangdong Second Provincial General Hospital for their technical assistance of live cell imaging system.

## Contributions

MHP and SCS designed the study. MHP performed the majority of the experiments. SML, ZNP, MHS, XHL, JQJ, YZ contributed to the regents and materials. MHP, XHO and SCS analyzed the data. MHP and SCS wrote the manuscript.

## Competing interests

There is no conflict of interest to declare.

## Ethics approval and consent to participate

Not applicable.

## Consent for publication

Not applicable.

## Funding

This work was supported by the National Natural Science Foundation of China (32170857); the National Key Research and Development Program of China (2021YFC2700100). There is no conflict of interest to declare.

## Abbreviations

FMNL2: Formin-like 2
FMNLs: Formin-likes
GV: germinal vesicle
GVBD: germinal vesicle breakdown
ATI: anaphase-telophase I
MI: metaphase I
MII: metaphase II
ER: Endoplasmic reticulum

## Notes

### Competing Interest Statement

The authors have declared no competing interest.

### Summary of Updates

Additional experiments were performed. The organization of panels of the figures were optimized. We also revised the manuscript according to the reviewers' comments.

